# Multiplex DNA fluorescence in situ hybridization to analyze maternal vs. paternal *C. elegans* chromosomes

**DOI:** 10.1101/2022.11.01.514763

**Authors:** Silvia Gutnik, Ahilya Sawh, Susan E. Mango

## Abstract

Recent advances in high-throughput microscopy have paved the way to study chromosome organization at the single-molecule level and have led to a better understanding of genome organization in space and time. During development, distinct maternal and paternal contributions ensure the formation of an embryo proper, yet little is known about the organization of chromosomes inherited from mothers versus fathers. To tackle this question, we have modified single-molecule chromosome tracing to distinguish between the chromosomes of two well-studied strains of *C. elegans* called Bristol and Hawai’ian. We find that chromosomes from these two strains have similar folding patterns in homozygous hermaphrodites. However, crosses between Bristol and Hawai’ian animals reveal that the paternal chromosome adopts the folding parameters of the maternal chromosome in embryos. This is accomplished by an increase in the polymer step size and decompaction of the chromosome. The data indicate that factors from the mother impact chromosome folding in trans. We also characterize the degree of intermixing between homologues within the chromosome territories. Sister chromosomes overlap frequently in *C. elegans* embryos, but pairing between homologues is rare, suggesting that transvection is unlikely to occur. This method constitutes a powerful tool to investigate chromosome architecture from mothers and fathers.

## Introduction

When an egg and sperm fuse to generate a new organism, not only genetic information in the form of the paternal and maternal genome is passed to the next generation. Oocytes and sperm contribute non-genetic factors, ranging from modified histones and DNA methylation to RNA and proteins loaded into oocytes and sperm (Xavier, Roman et al. 2019, Özdemir and Steiner 2021). These maternally and paternally contributed factors are crucial for the early stages of development and dictate translational or transcriptional regulatory steps (Conine, Moresco et al. 2013, Stoeckius, Grün et al. 2014, Robertson and Lin 2015). In addition, parental factors influence nuclear organization (Perino and Veenstra 2016, Hug and Vaquerizas 2018, Gentsch, Spruce et al. 2019, Woodhouse and Ashe 2020, Guo, Liu et al. 2021) For example, in *C. elegans*, modified histones H3K27me3, H3.3 and H3K4me2 are transmitted from parents to offspring (Katz, Edwards et al. 2009, Gaydos, Wang et al. 2014, Delaney, Mailler et al. 2018), while a host of chromatin factors influence the memory between generations by emerging mechanisms (Perez and Lehner 2019, Woodhouse and Ashe 2020, Burton and Greer 2021). A method to define the parental origin of embryonic chromosomes is needed to address the influence of these factors on genome organization.

Eukaryotic nuclei face a complex organizational problem, whereby 2 meters-long DNA is packaged into a confined space of 1-10μm in diameter. A large body of work has discovered layers of chromatin organization that include DNA loops, topological associating domains (TADs) and compartments (Misteli 2020). TADs reflect contiguous sequences that show physical interactions due to loop extrusion and range in size from 20 to 1000kb. They can define interactions between cis-regulatory regions and target promoters (Lupiáñez, Kraft et al. 2015, Batut, Bing et al. 2022, Huang, Seow et al. 2022) but seemingly at only some regions of the genome (Galouzis and Furlong 2022). Compartments are non-contiguous sequences that associate based on the transcriptional activity and histone modifications within those sequences (Wang, Su et al. 2016, Bian, Anderson et al. 2020), and in extreme examples, on cell type (Falk, Feodorova et al. 2019).

To define loops, TADs and compartments, many studies rely on sequencing-based methods that average the signal from thousands or millions of nuclei. Haplotype-resolved HI-C methods enable parental chromosomes to be distinguished within a diploid cell, but usually require averaging many chromosomes together (Li, Lin et al. 2021). To circumvent this difficulty, we have focused on chromosome tracing, which tracks chromosome and subchromosome organization at the level of single molecules. This method relies on reiterative DNA fluorescence in situ hybridization (FISH) for a molecular connect-the-dots approach (Wang, Su et al. 2016, Bouwman, Crosetto et al. 2022). Normally, chromosomes in diploid cells are distinguished by marking the two chromosome territories within the nucleus, but currently one cannot determine which chromosome is derived from sperm and which from oocytes. Here we extend our chromosome tracing method (Sawh and Mango 2020) to mark maternal and paternal chromosomes in *C. elegans*. Our strategy relies on F1 hybrid offspring from crosses of two closely related *C. elegans* strains, Bristol (N2) and Hawai’ian (HI), and utilizes their divergent genomic sequences to distinguish each chromosome territory. To analyze these hybrids accurately, we have designed and implemented strain-specific FISH probe sets, and developed an image analysis pipeline that accurately distinguishes the parental chromosomes in embryos. Using this new approach, we show a proof of principle by determining the chromosome conformation of wild-type maternal and paternal N2 and HI chromosome V. We define the degree of overlap between pairs of chromosomes and find that chromosomes intermingle frequently, but only rarely pair. In addition, we show that N2 and HI conformations are overall similar, opening up the possibility of using this approach for interrogation of inheritance effects.

## Results

### N2- and HI-specific probes selectively mark their respective chromosomes

To distinguish homologous but distinct chromosomes within a single nucleus, we focused on F1 hybrid offspring from crosses between divergent *C. elegans* strains (**Figure 1A**). We chose N2, the commonly used laboratory strain, and the related HI as crossing partners for four reasons: (1) the two strains can interbreed (2) they have been extensively characterized at the sequence level (Koch, van Luenen et al. 2000, Wicks, Yeh et al. 2001, Swan, Curtis et al. 2002, Thompson, Snoek et al. 2015); (3) HI is one of the most divergent *C. elegans* isolates from N2, with over 170 000 SNPs between the two strains as well as many insertions and deletions, which supplied regions across the two genomes to design strain-specific chromosome tracing probes (Andersen, Gerke et al. 2012, Thompson, Snoek et al. 2015, Crombie, Zdraljevic et al. 2019, Kim, Kim et al. 2019); (4) the two strains show a high level of synteny and are similar enough in sequence that a set of common chromosome tracing probes could be used to trace both N2 and HI chromosomes, keeping reagent costs low.

**Figure 1:**
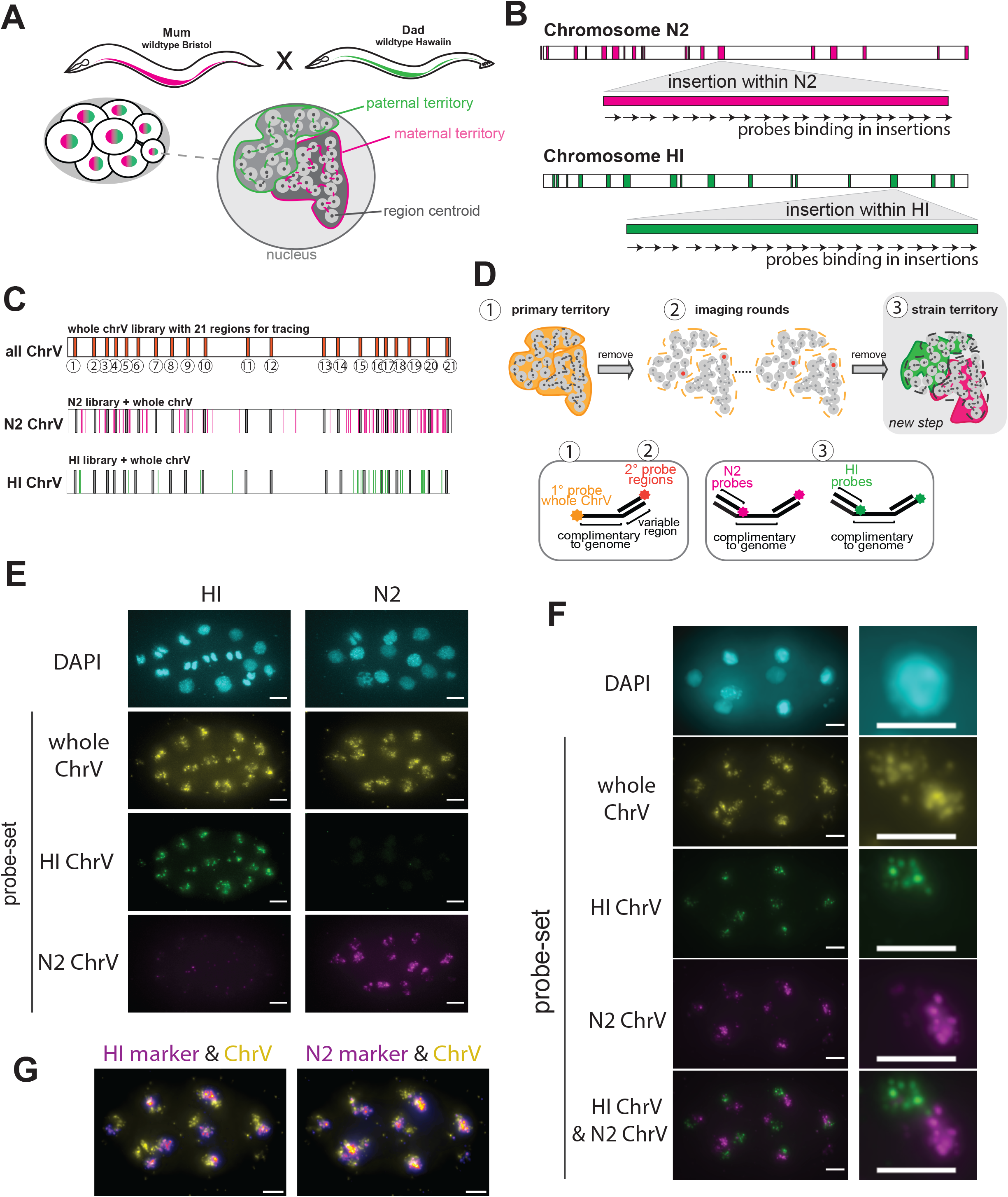
Haplotype-resolved chromosome tracing in *C.elegans*. (A) Schematic of crossing experiments (B) Schematic of probe design strategy (C) Location of whole chromosome V (ChrV) tracing library with 21 regions and N2 and HI libraries interspersed along ChrV. (D) Schematic representation of the imaging workflow using an automated imaging system (see methods), and probe design used for the indicated steps. (E) DNA Fish on embryos from N2 and HI, respectively, using the whole ChrV library, N2 ChrV and HI ChrV-marking libraries. Note that the N2 ChrV libraries causes a small punctual background staining in HI. Scale bar, 5μm. (F) DNA FISH on embryos derived from crosses between N2 hermaphrodites and HI males, using the whole ChrV library, N2 ChrV and HI ChrV-marking libraries. The haplotypes are well distinguishable. Scale bar, 5μm. (G) Overlay of ChrV territory signal with HI-marker (left) and N2 marker (right) in the embryo used in F. Scale bar, 5μm.

We made use of previously annotated insertions and deletions within the N2 and HI genomes (Thompson, Snoek et al. 2015) to design probes that were specific for each strain and could distinguish HI and N2 chromosome territories (**Figure 1B**). We focused on insertions larger then 1000 nucleotides (nts), which corresponded to a minimum of 33 potential 30-mer probes within a region, and which optimized the signal to noise ratio for DNA FISH. Across both genomes each chromosome harbored a varying number of sequence variants, from a high of 172 on ChrV to a low of 37 on the X chromosome (**Figure S1A**). The total length of inserted sequences larger than 1000 nts was the highest for ChrV (**Figure S1B**), and we therefore focused on ChrV to illustrate the proof of principle. We designed suitable probes to N2 and HI, as previously described (Rouillard, Zuker et al. 2003, Sawh, Shafer et al. 2020)(see Methods). This approach resulted in 5577 probes for N2 ChrV and 1831 for HI ChrV, with each probe set binding along the length of the respective chromosome, and located in-between the common tracing probes for ChrV (**Figure 1C & S1C**).

In conventional whole chromosome tracing of *C. elegans* embryos, fluorescently labeled primary DNA FISH probes are hybridized to defined regions along a chromosome such as the 22 regions along ChrV (Sawh, Shafer et al. 2020). When imaged *en masse*, the probes reveal the chromosome territory. (**Figure 1D panel 1**). Next, fluorescent region-specific probes are hybridized sequentially to readout tails on the primary probes, to visualize individual locations along ChrV (**Figure 1D panel 2**). We modified this method in two ways for our strain-specific approach. First, in addition to the common whole-chromosome tracing probes, we hybridized the strain-specific N2 and HI probes to the F1 hybrid embryos. In contrast to the common whole-chromosome tracing probes, we generated the strain-specific probes without a fluorophore, but with tails to bind two secondary oligos. These binding sites were unique for either the entire N2 or HI probe set, providing a means to label the strain territory by on-microscope hybridization during image acquisition. We note that restricting fluorophores to the secondary probes gave flexibility regarding the choice of fluorophore for each experiment and lowered the cost of probe synthesis. Second, we imaged the N2 and HI markers at the end of the chromosome tracing experiment (**Figure 1D panel 3**). We note, however, that since fluorescent labeling of the strain markers relies on on-stage hybridization during image acquisition, the strain markers could be imaged any time after primary probe imaging.

To test the specificity of the N2 and HI probes and assure compatibility of the new probe sets with the primary probe set for ChrV, we hybridized all three probe sets to fixed homozygous N2 and HI (CB4856) embryos. As expected, the primary probe set for ChrV enabled visualization of ChrV both in N2 and HI homozygous animals (**Figure 1E**). HI probes marked the HI chromosome without detecting a signal on N2. The N2 probes detected the N2 chromosome robustly. In addition, we observed a slight signal in the HI strain (**Figure 1E**). This signal likely corresponds to a small region within the HI genome that was mis-annotated in the Thompson genome, which we used for our probe design (Thompson, Snoek et al. 2015, Kim, Kim et al. 2019). Nevertheless, we found that this low signal did not interfere with image segmentation and chromosome classification, in subsequent experiments (see below). Future libraries could remove these sequences.

To assess if the strain-specific probe sets perform well with heterozygous embryos, we mated N2 mothers with HI fathers. The two chromosomes were clearly visible and N2 and HI probes marked their respective chromosome, allowing us to determine the parent of origin for each chromosome (**Figure 1F**). The N2 and HI signals overlapped partially with the chromosome V territory signal (**Figure 1G**), which reflects that the three probe sets target different sequences along ChrV. Together, it is clear which chromosome comes from which strain (**Figure 1F**). We conclude that the N2 and HI probe sets can distinguish ChrV derived from N2 or HI.

### N2 and HI chromosomes form barbells but HI is more compact

N2 and HI are both *C. elegans*, but they harbor sequence differences, some of which are predicted to affect chromatin architecture (Thompson, Edgley et al. 2013, Thompson, Snoek et al. 2015, Kim, Kim et al. 2019). For example, HI lacks the germline RNAi component *ppw-1*.Since experimentally introduced siRNAs were shown to be sufficient to induce chromatin compaction, it is possible that the lack of germline RNAi might influence chromosome conformation (Tijsterman, Okihara et al. 2002, Fields and Kennedy 2019). We therefore wanted to investigate if these two strains exhibited similar genome structures by comparing chromosomes from N2 or HI homozygous embryos. The newly derived N2 conformation was virtually identical to the previously published average N2 configuration for ChrV (Sawh, Shafer et al. 2020), with comparable step size (1.030 vs. 1.037) and scaling exponents (0.198 vs. 0.193) when fitted to a power-law function (**Figure 2A**). This result reveals the reproducibility of chromosome tracing in *C. elegans*, with little batch to batch variation between independent studies.

**Figure 2:**
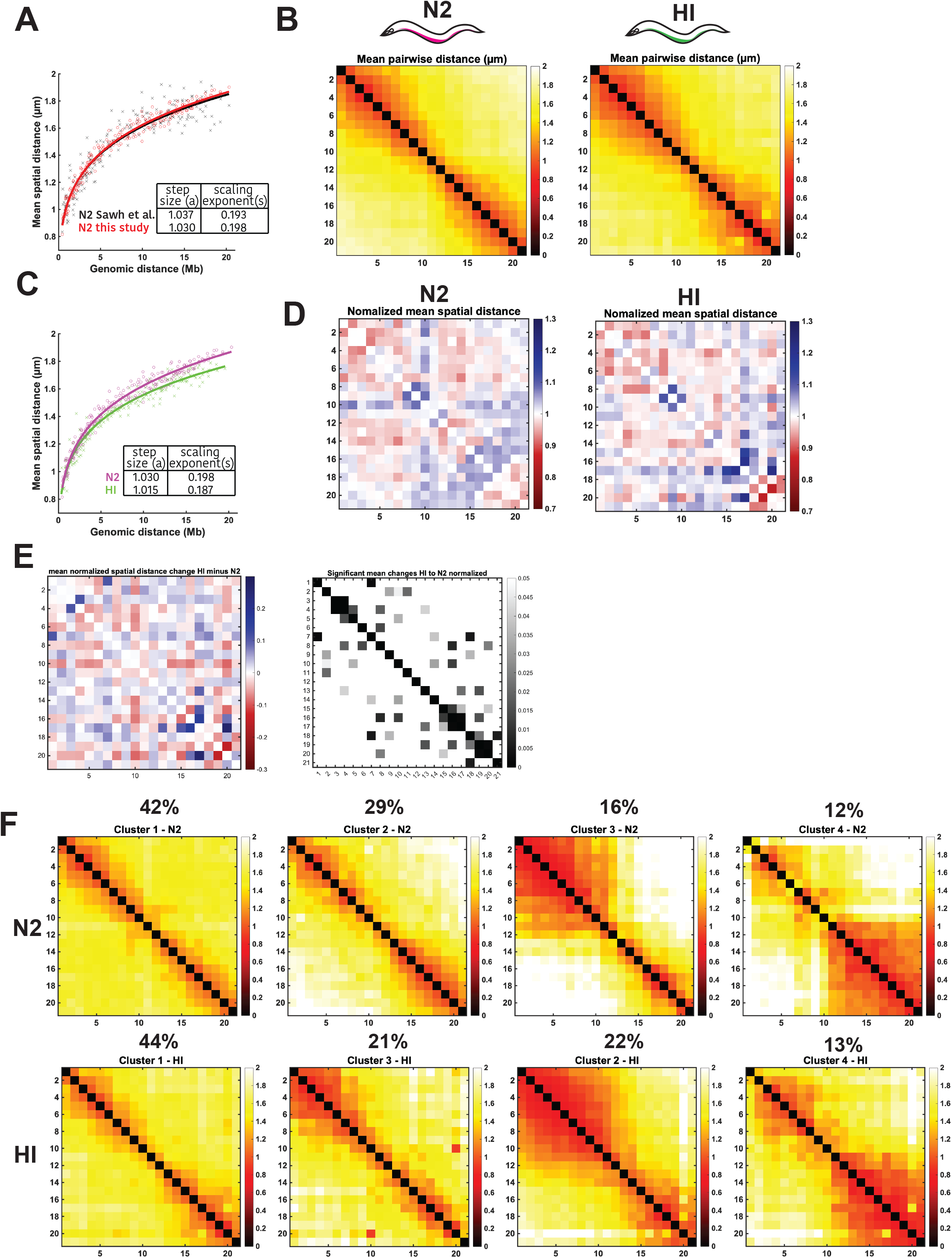
Highly similar N2 and HI ChrV organization. (A) Power-law fits of mean pairwise distances in μm versus genomic distance in Megabases (Mb) for N2 data generated in this study (black) and previously (red) (Sawh, Shafer et al. 2020). (B) ChrV mean distance matrix for N2 (left) and HI (right) in early embryos (2–40 cells), colored by distance (in μm). N = 6072 (N2) & N = 5905 (HI). (C) Power-law fits of mean pairwise distances for N2 data (magenta) and HI data (green). (D) Normalized spatial distances for N2 date (left) and HI data (right). Data is plotted as observed over expected. The expected spatial distance is determined by the fit to the powerlaw function shown in C. Red marks regions that are closer together than expected from the fit and blue regions that are further away. (E) Changes between normalized spatial distances of N2 and HI data (left) and p-value of the spatial distance changes. (F) Mean pairwise distances in μm of subpopulations of chromosome conformations for N2 and HI traces as determined by unsupervised clustering.

The HI chromosome showed similar overall properties to N2, with compacted chromosome arms and an extended center (**Figure 2B**). However, HI ChrV was slightly more compact than N2, as revealed by its smaller step size in power-law fitting (1.015 for HI vs. 1.030 for N2) (**Figure 2C**). N2 and HI also had differences in their scaling coefficients (0.198 for N2 and 0.187 for HI), which means that increasing genomic distances produces lower growth of spatial distances in HI compared to N2. These differences between N2 and HI may reflect the smaller size of the HI genome compared to that of N2 (Thompson, Snoek et al. 2015). They do not reflect differences in the diameters of interphase nuclei as we did not find a significant difference between N2 and HI nuclear sizes (**Figure S1D**).

Power-law fitting not only reveals folding properties of chromosomes, but it also serves as a mean to normalize the spatial distance measurements by taking into consideration the polymer nature of the chromosomes (Mirny 2011, Wang, Su et al. 2016, Sawh, Shafer et al. 2020). Normalization of the N2 and HI spatial distances revealed general similarities in folding complexities between HI and N2 (**Figure 2D**). For example, the distances within the left and right arms were smaller than predicted by the power law function for both N2 and HI, suggesting these regions are highly folded. In addition, the distances between the center and the right arm were larger than expected, consistent with an average barbell configuration for both strains (Sawh, Shafer et al. 2020). Despite the overall similarity of both strains, they showed some differences in some regional folding, as revealed by a number of differences in the normalized distances between N2 and HI (**Figure 2E**).

Previously, we had observed that individual chromosomes in vivo assumed conformations that were distinct from that of the population average (Sawh, Shafer et al. 2020). Similarly, when we performed unsupervised clustering on N2 and HI traces, we found the similar subpopulations of traces in both strains (**Figure 2F**). These consisted of chromosomes with one arm highly folded and chromosomes with both arms folded. Taken together, we conclude that N2 and HI chromosome V shows an overall similar structure to each other at the Megabase scale.

### A new pipeline to analyze N2:HI hybrids

Our next goal was to examine N2 and HI chromosomes after interbreeding. First, we developed and implemented a new image segmentation and tracing pipeline. Like conventional chromosome tracing, we applied watershed segmentation on the nuclear signal (DAPI-staining) to restrict the definition of chromosome territories to the nuclear volumes and remove background signals in the territory images that lay outside of the nuclei (**Figure 3A step 1**). Next, watershed segmentation was applied to images of chromosome territories. This step defined the volumes in which chromosomes were traced using a nearest-neighbor approach (**Figure 3A step 2; (Sawh, Shafer et al. 2020)**). In some instances (between 6-8% of traces), traces were ambiguous and excluded from further analysis. Third, strain-marking territories were segmented for N2 and HI, and the resulting volumes overlayed with the traces generated in the previous step (**Figure 3A step 3)**. Since the primary ChrV probes did not overlap perfectly with the strain marking probes, we classified traces into N2 or HI based on whether the trace was located closest to a strain marking volume for N2 or HI. We assessed the efficiency of this method to classify traces accurately by counting how often more than two N2 or HI traces were detected in one nucleus. We found only a minority of 2% of HI traces and 7% of N2 traces were mis-assigned and excluded these in downstream analysis.

**Figure 3:**
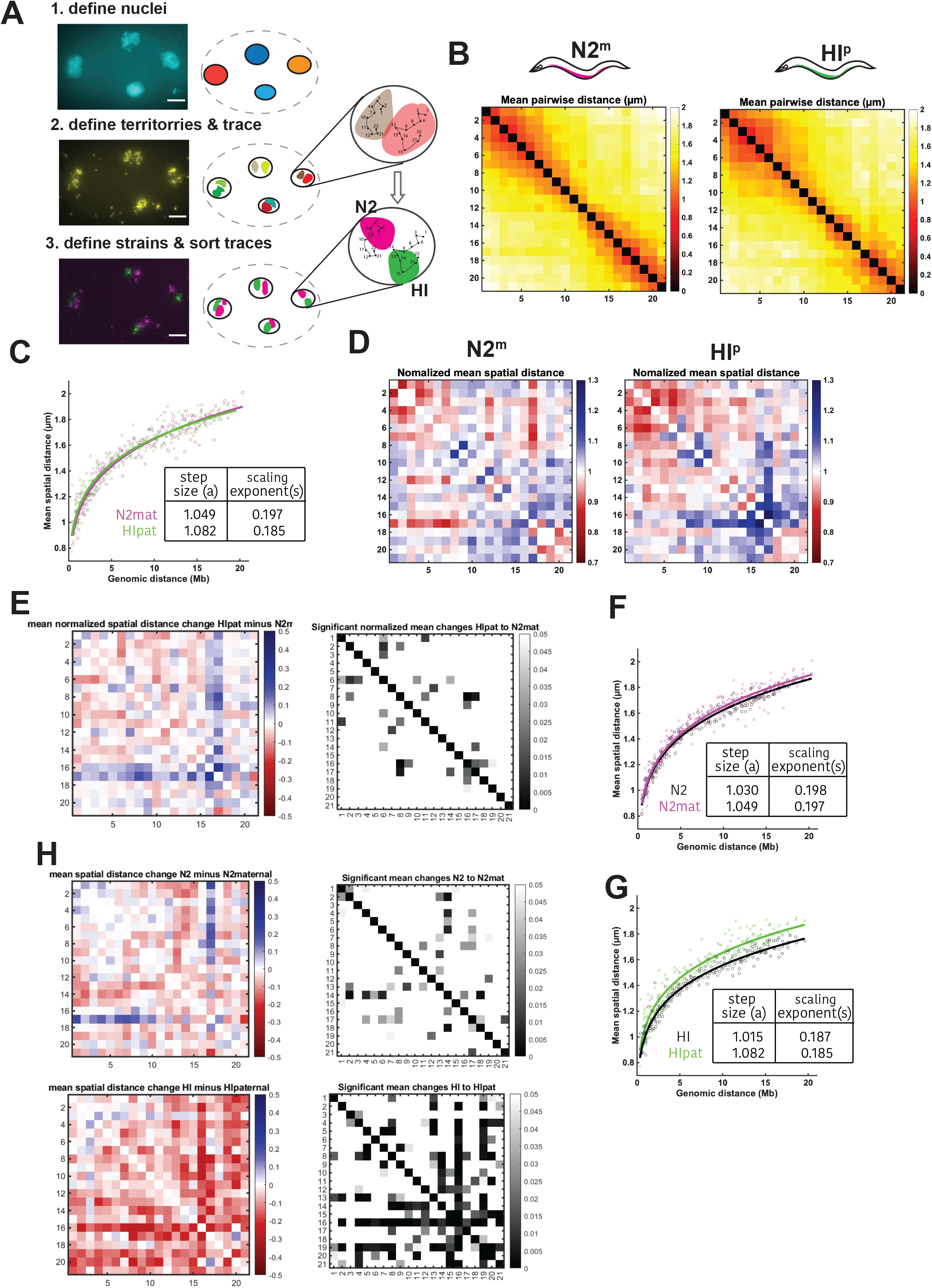
HI paternal chromosomes decompact when subjected to the N2 maternal environment. (A) Example of Z-projections of raw data collected during haploid-specific chromosome tracing and schematic representation of image segmentation, tracing and sorting of traces. Scale bar, 5μm. (B) ChrV mean distance matrix for traces derived from hybrid embryos (2-40cells) from crosses between N2 hermaphrodites and HI males, N2 maternal traces (left) and HI paternal traces (right), colored by distance (in μm). N = 1384 (N2^m^) & N = 1066 (HI^p^). (C) Power-law fits of mean pairwise distances for N2^m^ data (magenta) and HI^p^ data (green). (D) Normalized mean spatial distances for N2^m^ and HI^p^ traces shown as observed over expected spatial distances as determined by the power-law fit in C. (E) Differences in normalized mean spatial distances between N2^m^ and HI^p^ traces (left) and p-value (right). (F) Power-law fits of mean pairwise distances for N2^m^ data (magenta) and N2 homozygous data (black) (G) Power-law fits of mean pairwise distances for HI^p^ data (green) and HI homozygous data (black). (H) Differences of mean pairwise distances in μm (left), and p-value (right) of N2^m^ and N2 homozygous (top) and HI^p^ and HI homozygous (bottom)

### The paternal chromosome adopts the maternal conformation in hybrids

We used our new tracing pipeline on embryos derived from crosses between N2 hermaphrodites (N2^m^) and HI males (HI^p^). The overall conformation of ChrV in these hybrids agreed well with the homozygous conformations, with compacted arms and more open centers (**Figure 3B**). Power-law fitting revealed that both chromosomes showed almost identical genomic distances vs. spatial distance relationships with each other, and these resembled N2 homozygotes (**Figure 3C,F,G**). Only a minority of pair-wise distance changes were significant between N2 maternal (N2^m^) and HI paternal (HI^p^) chromosomes, indicating that they were highly similar overall (**Figure 3E**).

The N2^m^and HI^p^ chromosomes showed differences in step size (1.049 to 1.082) and scaling exponent (0.197 to 0.185). The scaling exponents were unchanged for HI and N2 chromosomes with respect to their homozygous conformations. The step size for HI^p^ chromosomes increased, from 1.015 to 1.082 in the N2^m^ background, whereas it remained virtually unchanged for N2^m^ compared to N2 homozygotes. Closer examination of mean pairwise distance changes of N2^m^ and HI^p^ chromosomes (compared to homozygous chromosomes from un-crossed embryos), revealed that substantially more regions changed in HI than in N2. Almost all these regions decompacted in HI^p^ chromosomes (**Figure 3H**). This result reveals that the HI^p^ chromosome decompacts when subjected to the N2^m^ environment and implies that the paternal chromosome is influenced by the maternal environment in trans.

To test the influence of the maternal HI (HI^m^) environment, we performed the reciprocal cross between HI hermaphrodites and N2 males. Again, we found that the overall conformation agreed well with HI and N2 homozygotes, with compacted arms and more open centers (**Figure 4AC**). Power-law fitting showed differences between N2 paternal (N2^p^) and HI^m^ chromosomes in the scaling exponent (0.200 to 0.164) and step size (1.069 to 1.131) (**Figure 4B**). As before, only a minority of N2^p^ and HI^m^ chromosomes pair-wise distance changes were significant (**Figure 4D**). This result confirms that HI and N2 chromosomes behave similarly within hybrids.

**Figure 4:**
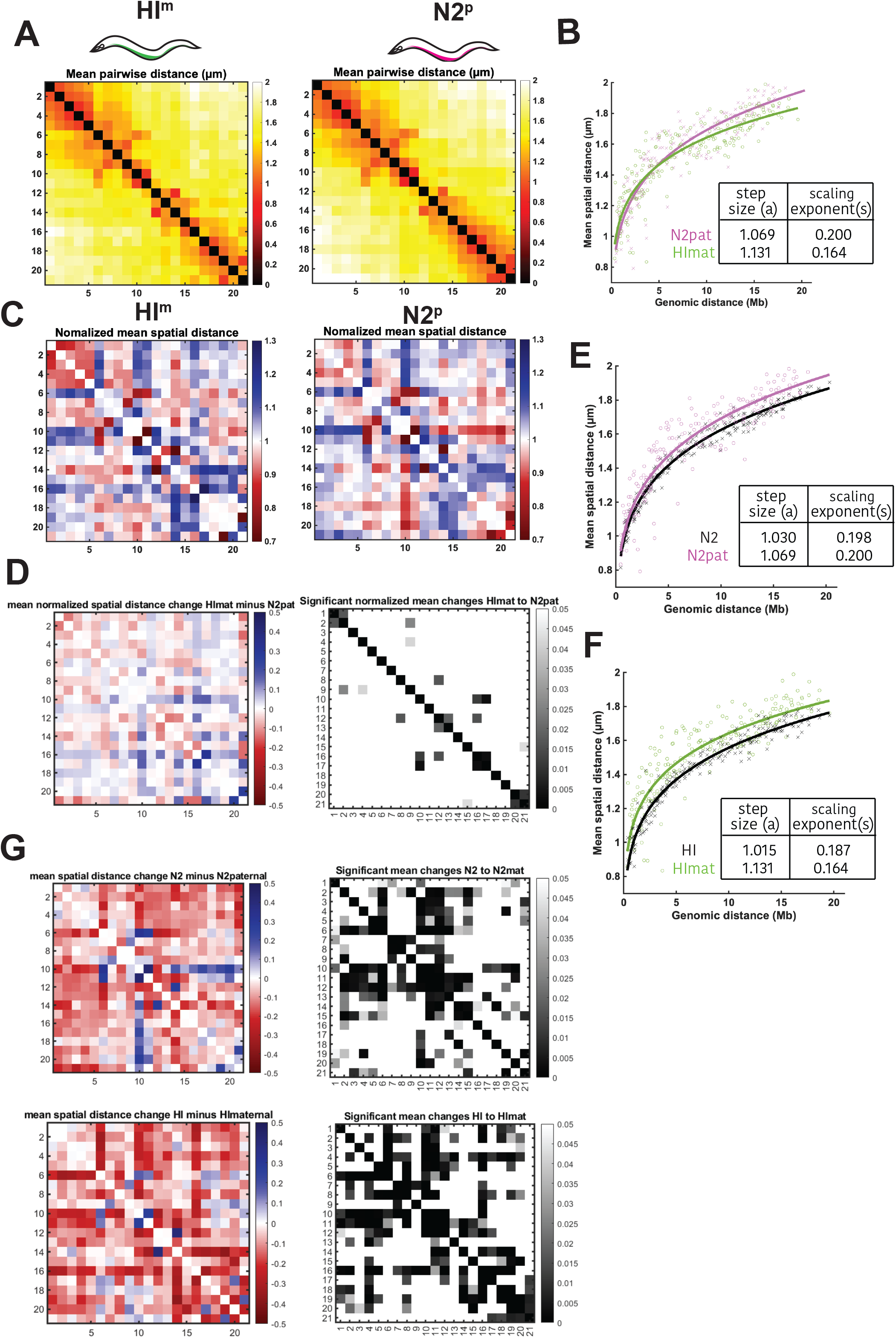
(A) ChrV mean distance matrix for traces derived from hybrid embryos (2-40cells) from crosses between HI hermaphrodites and N2 males, HI maternal traces (left) and N2 paternal traces (right), colored by distance (in μm). N = 1125 (HI^m^) & N = 1254 (N2^p^). (B) Power-law fits of mean pairwise distances for N2^p^ data (magenta) and HI^m^ data (green). (C) Normalized mean spatial distances for N2^p^ and HI^m^ traces shown as observed over expected spatial distances as determined by the power-law fit in B. (D) Differences in normalized mean spatial distances between N2^p^ and HI^m^ traces (left) and p-value (right). (E) Power-law fits of mean pairwise distances for N2^p^ data (magenta) and N2 homozygous data (black) (F) Power-law fits of mean pairwise distances for HI^m^ data (green) and HI homozygous data (black) (G) Differences of mean pairwise distances in μm (left), and p-value (right) of N2^p^ and N2 homozygous (top) and HI^m^ and HI homozygous (bottom)

The paternal (N2) chromosome decompacted compared to the homozygous (N2) chromosome as indicated by the increased step size (1.069 vs. 1.030) and constant scaling exponent (0.20 to 0.198)(**Figure 4E**). Interestingly the HI^m^ chromosome changed in two ways compared to the HI homozygous chromosome, first by a decreased scaling exponent and second by an increased step size (**Figure 4F**). This result suggests that each chromosome is subtly changed in this particular hybrid (**Figure 4E,F**). When we compared mean pair-wise distance changes of HI^m^ and N2^p^ chromosomes with their homozygous counterpart, we detected significant changes for both chromosomes. Consistent with the decompaction we found by power-law fitting, most regions became decompacted.

Taken together these data suggest that paternal chromosomes are influenced by the maternal environment irrespective of the strain. Since we also find that HI^m^ chromosomes change in the presence of N2^p^, we hypothesize that the N2 chromosomes influence HI chromosomes in trans, while N2 chromosome structure seems to be resistant to influences by the HI chromosome.

Previous data showed that averaging many traces could mask distinct subpopulations of conformations that exist in vivo (Sawh, Shafer et al. 2020). We wondered if closer examination of chromosome conformation might reveal differences between paternal and maternal chromosomes. To answer this question, we performed unsupervised clustering (Sawh, Shafer et al. 2020) on N2^m^ and N2^p^, as well as HI^m^ and HI^p^ chromosome traces from 2-40 cell stage embryos. We found that the dominant clusters with a contribution of 36% and 28% (N2^m^), and 38% and 25% respectively (HI^m^) had low levels of folding in the arms separated from each other (**Figure 5A panels 1 & 2 and Figure 5B panels 1 & 2**). In addition, N2^m^ chromosomes had clusters with either a highly folded right or left arm, as seen previously, whereas HI^m^ only had a highly folded left arm. In addition, 18% of HI^m^ traces showed a conformation with a compacted central domain and arms close to each other (**Figure 5B panel 4**). Taken together, maternal traces from both strains were similar in the subpopulations of conformations. In contrast, N2 and HI paternal traces had bigger differences (**Figure 5CD**). While HI^p^ subpopulations were characterized by folding of one or the other chromosome arm, N2^p^ clusters were more open and a subpopulation with a highly folded right arm was not present.

**Figure 5:**
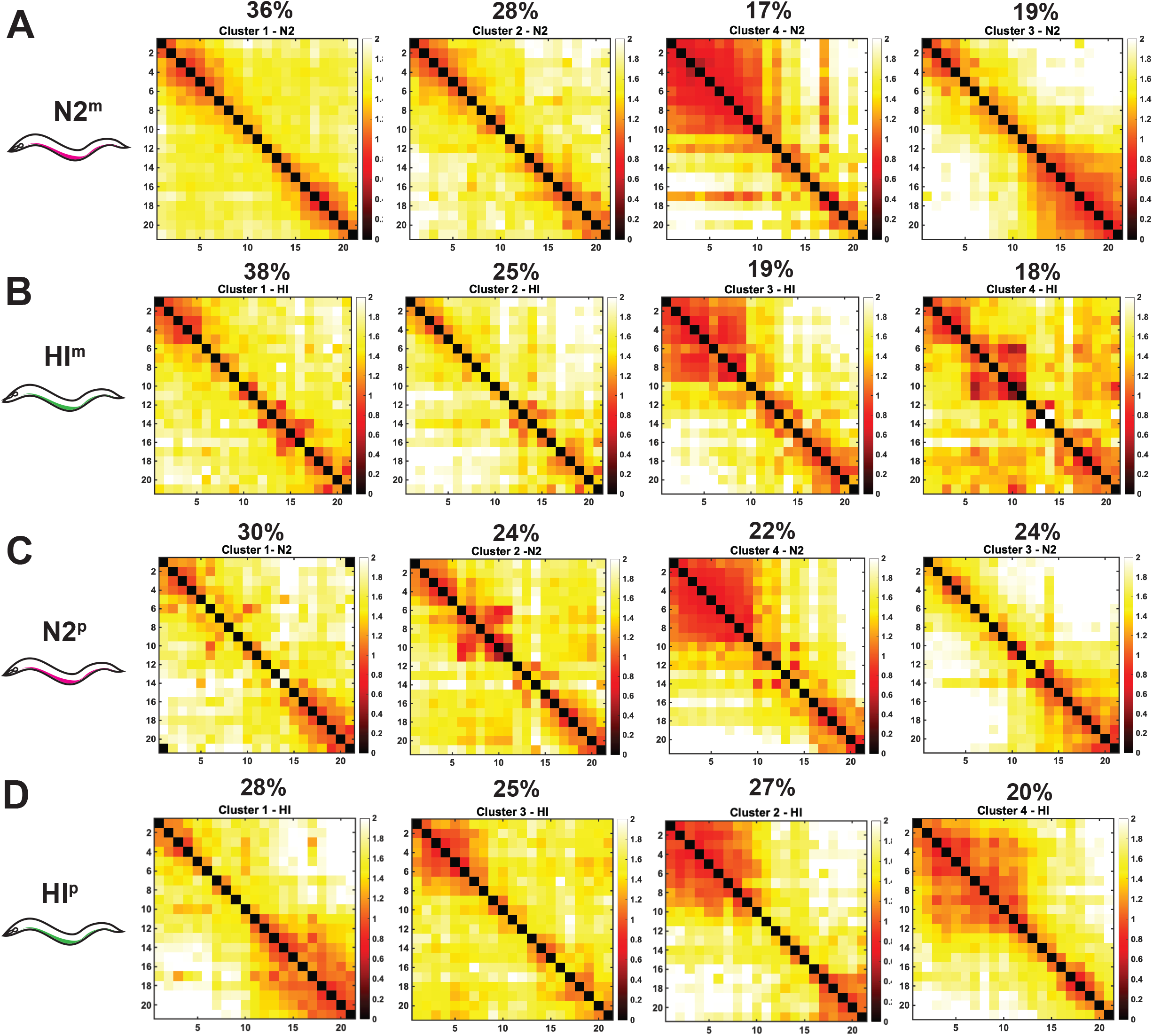
Mean pairwise distances in μm of subpopulations of chromosome conformations for N2^m^ (A) HI^m^ (B), N2^p^ (C) and HI^p^ (D), ordered by similarity to subpopulations in (A).

### Homologous chromosomes do not align

Transcription regulation depends on cis-regulatory sequences that are typically adjacent to target promoters (Robson, Ringel et al. 2019, Huang, Seow et al. 2022). In *Drosophila* and other Dipterans, homologous chromosomes are paired in somatic cells, allowing for interchromosomal interactions between enhancers and promoters, a phenomenon termed transvection (McKee 2004). Methods to study physical interactions of homologous chromosomes are limited since the distinction between homologous chromosomes is not always possible (Li, Lin et al. 2021).

To determine the degree of overlap between homologs within single nuclei, we used stringent image segmentation on Z-stacks of the N2 and HI markers acquired during tracing of N2^m^/HI^p^ and HI^m^/N2^p^ hybrid embryos. We defined the territory volume the N2 and HI ChrV occupied as the number of voxels which contained a signal for N2 or HI. The overlap was then calculated as the number of voxels which were marked by N2 and HI divided by total number of voxels occupied by N2 or HI marking (**Figure 6A**). Since we detected a small background staining in N2 embryos with the probes that are designed to mark the HI strain (**Figure 1E**), we used a cut-off of 5%. Below this we considered the homologs non-overlapping. We found that 37% of nuclei showed no overlap and of the remaining nuclei, 43% had an overlap larger than 15% of the strain-marking volume. Thus, sister chromosomes display frequent territory overlap in *C. elegans* embryos.

**Figure 6:**
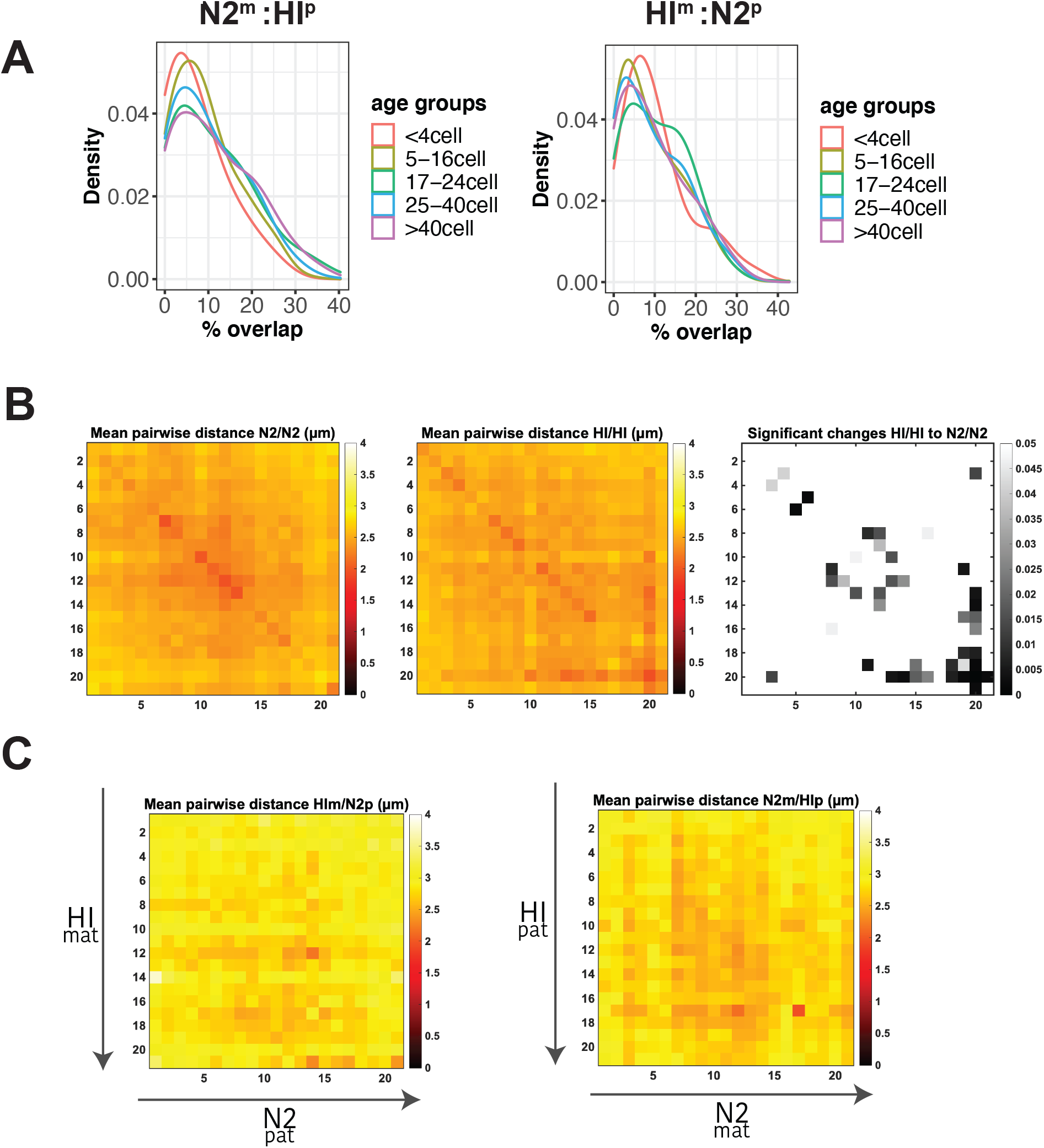
(A) Density map of overlaps between N2 and HI chromosomes from crosses between N2^m^ and HI^p^ in different age-groups of embryos indicated by color-coding. The % overlap is defined as the ratio between voxels of overlapping N2 and HI marker signal, and total number of voxels defining the N2 territory and HI territory. (B) Mean spatial distances between all regions of one homolog to all regions of the other homolog of N2 (left) and HI (middle) ChrV in homozygous embryos, colored by distance (in μm). (C) Mean spatial distances between all regions of one homolog to all regions of the other homolog of HI^m^ and N2^p^ chromosomes (left) and of HI^p^ and N2^m^ (right), colored by distance (in μm).

We next asked if certain regions along sister chromosomes were preferentially overlapped. Chromosome tracing records the trajectory and hence the location of individual regions of single chromosomes within single nuclei. To determine whether individual regions of one homolog were preferentially closer to individual regions on the other homolog, we first focused on traces derived from homozygous N2 and HI embryos, respectively. We filtered our data to restrict the analysis to all nuclei that contained exactly two traces and therefore had not yet undergone DNA replication. We measured the distances of all regions along one chromosome to all regions along the homologous partner chromosome within the same nucleus for N2 and HI homozygous embryos (**Figure 6B**). Distances were generally fairly large, on the order of 2.5 μm in nuclei that are on average 5.3 μm in diameter. While this absolute distance may be skewed by fixation conditions, this result suggests that although sister chromosomes frequently overlap a portion of their territories, they are not closely aligned with one another. We also note that there were few significant differences for chromosomes from N2 vs HI.

We also examined our hybrids and measured pair-wise distances between all regions along HI^m^ and N2^p^ chromosomes, and N2^m^ and HI^p^. We restricted the analysis again to all nuclei that contained exactly two traces. The data revealed that *C. elegans* does not stably align homologous domains and inter-homolog interactions might, similar to mammalian genomes, be confined to particular regions of the chromosome and occurring within specific cell types (Apte and Meller 2012). Interestingly, we observed that distances between HI^m^/N2^p^ chromosomes were typically larger than those for N2^m^/HI^p^.

## Discussion

This study analyzes chromosome conformations from maternal and paternal genomes using crosses between divergent *C. elegans* strains. We have developed probes and an analysis pipeline to distinguish between N2 and HI strains after FISH, and we have used these tools to determine the conformations of maternal vs. paternal ChrV at the Megabase scale. To our knowledge this is the first adaptation of chromosome tracing to distinguish maternal vs. paternal chromosome conformations during development.

Our data reveal that i) N2 and HI embryos have similar chromosome conformations, with few differences in particular regions; ii) after inter-strain crosses, HI chromosomes adopt the configuration of the N2 maternal configuration, implicating maternal factors that act in trans; iii) sister chromosomes frequently intermingle their territories, however homologous alleles rarely align.

Studies of haploid-resolved chromosome conformations have been largely limited to Hi-C methods, which tend to rely on population averaging. Our tracing and territory approach enabled us to determine the conformation of many regions along a chromosome, not only those with SNPs or indels. In this first study, the strain marking library relied on insertions larger than 1000nts. Future libraries could increase the density of probes by including sequences from smaller insertions. This adaptation would in turn allow the discrimination of less divergent strains or, in the case of tracing libraries for subregions of chromosomes, the marking of chromosomes and regions which are less divergent. The approach we presented therefore is adaptable for usage in a variety of conditions. In addition, chromosome tracing by multiplexed FISH in general has the added benefit of preserving the spatial properties of the region under study within the nucleus, such as proximity to the nuclear periphery. It also preserves the tissue in study, making it possible to relate chromosome traces to the presence of cellular features outside of the nuclei (Wang, Su et al. 2016, Bintu, Mateo et al. 2018, Liu, Lu et al. 2020, Sawh, Shafer et al. 2020).

Our data have revealed that the chromosome conformations of HI and N2 are similar and resemble barbells, as seen previously for N2 homozygotes (Sawh, Shafer et al. 2020). This is true for both self-progeny and cross-progeny between strains. A minority of pairwise-distances along ChrV showed significant differences in inter-probe distances, even though ChrV is the most divergent of chromosomes between the strains. Long-read sequencing had identified several large rearrangements, one of which translocated 170kb of N2 ChrV left arm to the left arm of ChrII in CB4856 (Kim, Kim et al. 2019). None of these rearrangements or SNPs influenced the larger-scale chromosome organization of ChrV. The clusters of subpopulations between the two strains were similar as well. We observed two clusters with one of the two chromosome arms being compacted by long-range folding, and two clusters with more uniform compaction along the entire chromosome. Despite their average similarity, HI clusters showed smaller distances in left-right-arm measurements compared to N2, suggesting the inter-arm proximity may account for the higher compaction of HI ChrV compared to N2 ChrV.

Using our haploid chromosome tracing method, we found that paternal traces are overall less compacted than the homozygous population of traces of the same strain. One hypothesis is that paternal traces are subjected to a bigger influence by the maternal environment. Comparisons of our traces show few regional differences, which might reflect the robustness of the larger scale chromosome organization. We note that the resolution of the library was not high enough to detect smaller, regional differences between alleles. Therefore, we cannot exclude that maternal and paternal chromosomes are different on the local chromatin level. Future studies with smaller-scaled chromosome tracing libraries are needed to tackle this question.

In Drosophila and other Dipterans, homologous chromosomes are paired in somatic cells (McKee 2004). Up until now, strict pairing of homologs in other organisms has not been observed. In mammals pairing of homologs is tissue specific and restricted to particular regions of chromosomes (Apte and Meller 2012). Earlier studies in mice using FISH demonstrated that chromosomes tend to locate in heterologous neighborhoods (Khalil, Grant et al. 2007) and work in human cells interestingly found larger distances between homologs then heterologs (Heride, Ricoul et al. 2010). These observations imply that inter-homolog interactions may be rare. In this study, we found that in *C. elegans* homologs of ChrV frequently intermingle, but homologous regions rarely closely align.

Among all embryos scored for overlapping sister chromosomes, the transcriptionally inactive 2-4 cell stage embryos displayed the least. After the first zygotic genome activation (ZGA, 4-cell stage), embryos exhibited an increase in sister overlap (Figure 6A, arrow). In other species, chromosomal regions of actively transcribed genes tend to localize together (Branco and Pombo 2006), suggesting that transcription might play a role in the increased interactions between homologs in *C. elegans* as well. Since we see an increase in overlap between homologs right after ZGA, when nuclear sizes are not very different (5-16cell), it is unlikely that decreased nuclear size is a driver of intermingling. An alternative hypothesis for increased chromosomal interaction in these very early stages could be differences in the local epigenetic landscape of very early embryos. Constitutive heterochromatin in the form of H3K9 methylation accumulates gradually in early *C. elegans* embryos (Mutlu, Chen et al. 2018). Thus, it is possible that the increased territory intermingling and heterochromatin formation are linked phenomena.

## Materials and Methods

### Strain maintenance and crossing experiments

Bristol (N2) and Hawai’ian (CB4856) strains were maintained at 20°C and grown on OP50 (Brenner 1974). For crossing experiments, the evening before embryo collection mid/late L4 hermaphrodites were placed together with equal number of L4 or young adult males of the opposite strain on a mating plate (regular 6cm plate with agar cut to ~1/6 size).

### Probe design and synthesis

#### Strain-specific probes set

Pools of oligo probes were generated from sequences that encompassed insertions > 1000nts for ChrV of Bristol (N2) and Hawai’ian HI (CB4856; (Thompson, Snoek et al. 2015)). For each probe region, we extracted the genomic sequence of the *C. elegans* genome assembly (Ce10 for Bristol and Thompson for HI). The probes were predicted by OligoArray2.1 (Rouillard, Zuker et al. 2003), which uses NCBI-BLAST 2.2.26 and the OligoArrayAux secondary structure predictor. The parameters used were: melting temperature 60-100°C, no cross-hybridization or predicted secondary structure with a melting temperature greater than 70°C, GC content 30%–90%, no stretches of 7 or more identical nucleotides. OligoArray2.1 was run on the SCICORE high performance computing cluster at the University of Basel.

Next, the predicted 30nt probes were checked for unique sequence binding to the genome of interest using NCBI BLAST 2.9.0+ (Ce10 for N2-specific probes and Thompson-genome for HI-specific probes). In addition, all probe sequences were confirmed to not bind to the other genome by NCBI BLAST 2.9.0+ (Thompson for N2-specific probes and Ce10 for HI-specific probes). Probes sequences were then fused to tail sequences that included primer binding sites for amplification and a secondary oligo binding sites unique for either the entire N2-specific or the HI-specific probe-sets.

#### Whole-Chromosome tracing probes

Whole Chromosome tracing probes for ChrV were as described previously, with 22 100kb regions located in the center of previously identified TADs (Sawh, Shafer et al. 2020). Due to low signal to noise ratio for one of the 22 regions of this tracing library we excluded it from our experiments.

The probe libraries were amplified and labeled with fluorophores (whole chromosome library only) (Wang, Su et al. 2016, Sawh and Mango 2020, Sawh, Shafer et al. 2020). In brief, probes were amplified using limited cycle PCR (Phusion Hot Start Master Mix, Life Technologies), high yield in vitro transcription (HiScribe T7 Quick High Yield RNA Synthesis Kit, NEB), and cDNA synthesis reactions (Maxima H Minus Reverse Transcriptase, Fisher Scientific). Only for the whole Chromosome library (ChrV), probes were 5’ labeled with a fluorophore (ATTO 565 (IDT)) during the reverse transcription step. Probes were purified using: the DNA Clean & Concentrator kit with Spin IC columns after PCR, and Spin V columns using Oligo Binding Buffer after cDNA synthesis (all reagents from Zymo).

### Probe amplification

#### Embryo sample preparation and in situ hybridization

Embryos were dissected, fixed and hybridized to DNA FISH probes as described previously (Sawh, Shafer et al. 2020). In brief, embryos from young gravid adults were dissected in ddH2O and transferred to a round cover-slip (Bioptechs) coated with Poly-L-Lysine (Sigma) where). Embryos were fixed with 1% paraformaldehyde/ 0.05% Triton X-100, frozen on dry ice and freeze cracked (Kiefer, Smith et al. 2007, Sawh and Mango 2020). After washing once with 1x PBS and three times with 1xPBS/0.5% Triton, samples were treated with RNaseA (0.05mg/ml) for 30min at 37°C and blocked using hybridization buffer (10% dextran sulfate / 0.1% Tween-20/ 2X SSC/ 50% formamide) for 1hr at 37°C. Primary probes for the whole chromosome library (ChrV) and strain-specific probe sets were diluted to ~1μM each in hybridization buffer. Probes were annealed at 80°C on a hot metal plate for 10min and hybridized for at least 16hrs at 37°C. Slides were first washed with 2XSSC/50% formamide for 1hr at 37°C and washed additionally 2-times using 2xSCC and 2-times 0.5x SCC using prewarmed buffers. Slides were then stored at 4°C or imaged immediately.

#### DAPI staining and microscopy chamber assembly

Nuclear staining and flow chamber assembly were performed as described previously (Sawh and Mango 2020). Fixed embryos on round cover slips were stained with DAPI in 2xSCC (1:1000, ThermoFisher) for 5-10min immediately prior to imaging and washed 3-times with 2xSCC for 5-10min.

Specimens were mounted using a flow cell (Focht Chamber System 2 (FCS2®), Bioptechs), equipped with a micro aqueduct and attached to a home-build fluidic system (Moffitt and Zhuang 2016) attached to the microscope. During flow cell assembly 0.1μm Tetraspeck (ThermoFisher) beads in 2xSCC were allowed to adhere to the coverslip in order to track sample drift during image acquisition.

#### Image acquisition

Images were taken on a Nikon Ti2 equipped with a Photometrics Prime 95B camera, Lumencor SpectraX light source and Omicron lasers for bleaching the primary probe signals. Sequential hybridizations, image acquisition and bleaching of probe signals were operated by an automated program implemented in the NIS Elements Software as described previously (Sawh, Shafer et al. 2020). At the start of an acquisition embryos were selected by strong primary probe signal. For each FOV the first round of imaging consisted of primary probe imaging using 561nm illumination, nuclear stain imaging using 405nm illumination and fiducial bead imaging using 488nm illumination, and a total of 30μm were acquired in Z using 200nm steps for all imaging rounds. The primary probe signal was then bleached to undetectable levels using the Omicron lasers. The following sequential secondary probe imaging steps consistent of 20min incubation of secondary probes (8nm) on stage, washing in 2XSSC/25% Ethylene carbonate and imaging using 561nm, 647nm illumination (for probe signals) and 488nm illumination (for bead signals), followed by photobleaching. These steps were repeated for all secondary probe regions sequentially. For acquisition of strain-specific markings an additional hybridization and imaging step was added, which consisted of 20min incubation of secondary probes (32nm), washing in 2XSCC/25% Ethylene carbonate and imaging using 561nm,647nm and 488nm illumination.

### Data analysis

#### Generation of chromosome traces

Foci fitting and assignment was done as published previously (Wang, Su et al. 2016, Sawh, Shafer et al. 2020). Image segmentation and chromosome tracing was done in MATLAB as described previously with modifications (Sawh, Shafer et al. 2020). In brief, for each embryo separately, first nuclei were identified using the DAPI signals. Background signal was subtracted, then noise removed and strong signal smoothened. The image was binarized and the distance transform was computed on the resulting binary image prior to watershed segmentation. This resulted in a volumetric mask of individual nuclei. Next territories were segmented analogous to nuclei segmentation, whereby signal outside of nuclei were excluded using the nuclear mask. This resulted in a volumetric mask, where each volume represented a chromosome. Tracing of chromosomes was then performed within the defined chromosome volumes as done previously using the nearest-neighbor approach.

#### Assignment of traces to N2 or HI in crossing experiments

For experiments where strains were marked using the N2 and HI ChrV library, the strain marking signals were segmented as done for territories. The generated traces were then classified into HI and N2 traces based on whether the majority of the foci detected were located within or closest to a strain marking territory. Traces shorter than 4 regions were excluded to prevent misassignments and ambiguous traces, where equal number of N2 and HI was identified, were removed.

#### Normalization of distances by power-law fitting and statistical analysis

Spatial distance normalization and statistical analysis of distances was performed as described previously (Wang, Su et al. 2016, Sawh, Shafer et al. 2020).

#### Cluster analysis

Unsupervised clustering was performed as described previously (Sawh, Shafer et al. 2020). Since Region 17 resulted in fewer datapoints detected in tracing experiments, it was excluded from Cluster analysis. Resolutions used were as follows: 0.7 for HI traces, 0.6 for N2 traces, 1.0 for Hi^p^ and N2^m^, 0.7 for Hi^m^ and 0.9 for N2^p^ traces.

#### Analysis of homolog overlap

To calculate the overlap between N2 and HI chromosome territories, stringent image segmentation, as described above, was applied on a subset of images taken for chromosome tracing of N2^m^:HI^p^ and HI^m^:N2^p^ embryos. The generated 3D masks of N2 and HI markers were overlayed, and the voxels counted where N2 and HI overlapped. The %-overlap was defined as the ratio between overlapping voxels and the total voxel count of N2 and HI marker.

#### Intra-homolog distance measurements

Data generated in tracing experiments on N2 homozygous, HI homozygous and N2^m^:HI^p^ and HI^m^:N2^p^ embryos was used to measure distances between all regions of one homolog to all regions of the other homolog, analogous to inter-chromosome distance measurements. Data was filtered for traces which had exactly one partner trace within the same nucleus. Since watershed segmentation occasionally failed to segment nuclei well, partner traces which were found to be further then 6μm away between any region, were exclude, since these most likely resulted from mis-segmentation. The distances of all regions on one homolog to all regions of the other homolog was then calculated on the remaining partner chromosomes.

---

**Figure S1.**
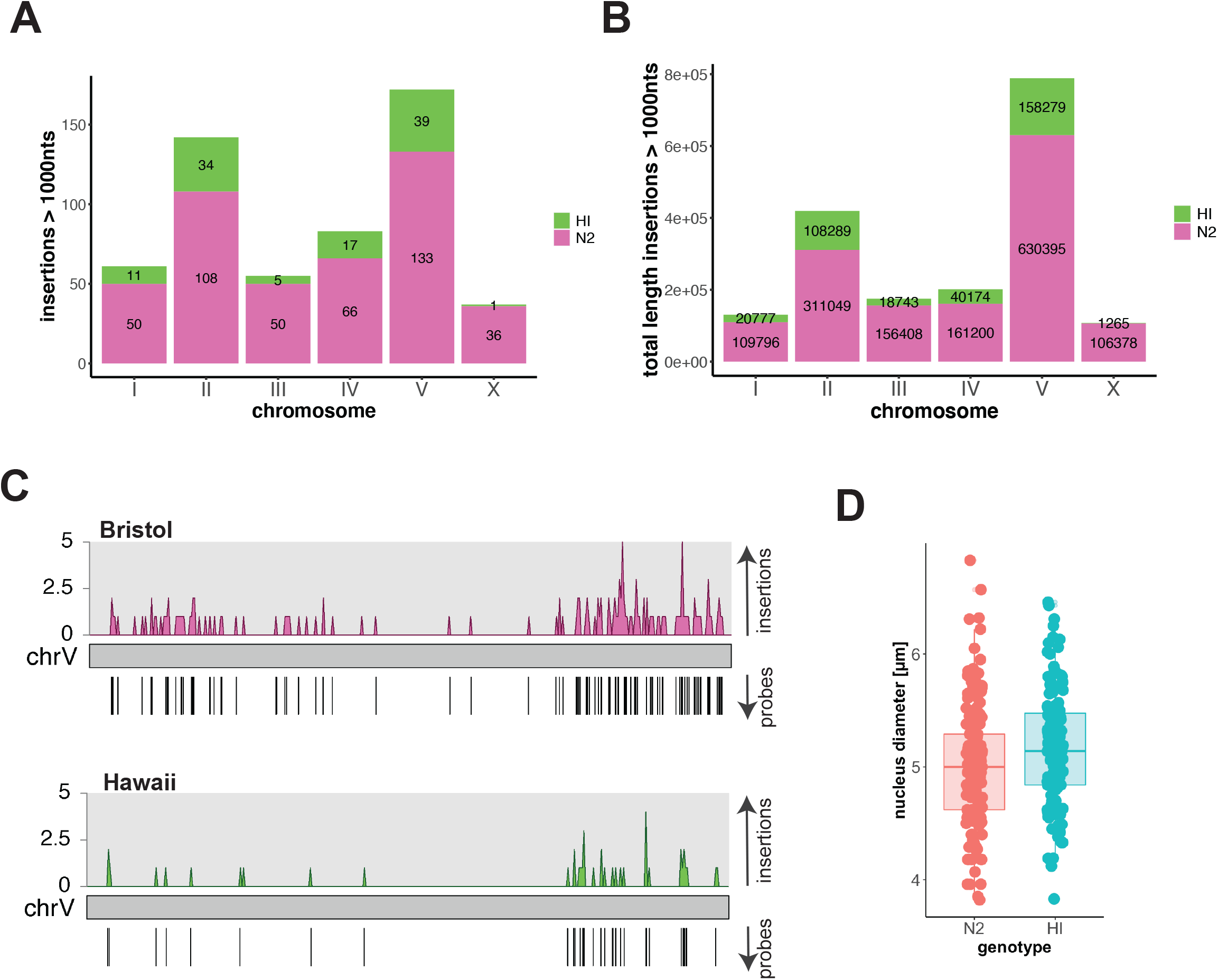
(A) Number of insertions larger then 1000nts present on chromosomes of N2 (magenta) and HI (green). (B) Total length of insertion larger then 1000nts present on chromosomes of N2 (magenta) and HI (green). (C) Location-density of insertions along N2 (magenta) and HI (green) ChrV binned at 50.000nts (top), and location of final strain-marking probes (bottom). (D) Nuclear diameter in N2 (red) and HI (blue) 4-8cell stage embryos.

